# Deletion of intestinal SHP impairs short-term response to cholic acid challenge in mice

**DOI:** 10.1101/2020.09.02.280230

**Authors:** James T Nguyen, Ryan R Riessen, Tongyu Zhang, Colin Kieffer, Sayeepriyadarshini Anakk

## Abstract

Small heterodimer partner (SHP) is a crucial regulator of bile acid (BA) transport and synthesis; however, its intestine-specific role is not fully understood. Here, we report that Intestine-specific *Shp* knockout (*IShpKO*) mice have higher intestinal and hepatic BAs, but not serum BAs when challenged with an acute (5-day) 1% cholic acid (CA) diet. Consistent with this finding, BA synthetic genes *Cyp7A1* and *Cyp8b1* are not repressed to the same extent in *IShpKO* compared to control mice post-CA challenge. Loss of intestinal SHP did not alter *Fxrα* mRNA but increased *Asbt* (BA ileal uptake transporter) and *Ostα* (BA ileal efflux transporter) expression even under chow-fed conditions. Surprisingly, the acute CA diet in *IShpKO* did not elicit the expected induction of *Fgf15* but was able to maintain the suppression of *Asbt*, and *Ostα/β* mRNA levels. At the protein level, ASBT was downregulated, while OSTα/β expression was induced and maintained regardless of diet. Examination of ileal histology in *IShpKO* mice challenged with acute CA diet revealed reduced villus length and goblet cell numbers. However, no difference in goblet cell number, villus morphology, crypt depth, and the expression of BA regulator and transporter genes was seen between *f/f Shp* and *IShpKO* mice after chronic (14-day) CA diet suggesting an adaptive response. We found the upregulation of the *Pparα*-*Ugt* axis, which can reduce the BA burden and compensate for the ileal SHP function. Thus, our study reveals that ileal SHP expression contributes to both overall intestinal structure and BA homeostasis.

## Introduction

Bile acids (BAs) are biological surfactants that allow for the digestion of lipids and fats (1,2). Over the last two decades, their roles in regulating various metabolic processes involved in health and disease have been uncovered. For example, BAs also play major roles in glucose, fat, and lipid metabolism and energy expenditure (3–5). BAs are synthesized from cholesterol and then secreted into the intestine to aid in fat digestion and 95% are recycled back to the liver. Defect in BAs recycling leads to lipid malabsorption and intestinal inflammation (6). Thus BAs concentrations are maintained by negative feedback coordinated by nuclear receptors (7), farnesoid X receptor (FXR), and small heterodimer partner (SHP).

FXR is an endogenous BA receptor and the BA-FXR-SHP axis is known to modulate BA synthesis, hydrophobicity, and transport (8). *Fxr-null* mice exhibit increased serum BA levels in response to cholic acid (CA) diet due to the loss of this negative feedback. When placed on a high cholesterol diet, *Fxr-null* and liver-specific *Fxr* knockout exhibited an increase in hepatic lipid levels, whereas intestine-specific *Fxr* knockout display lower hepatic lipid levels (9,10). Similarly, both liver-specific and intestine-specific *Fxr* knockout show deficits in the suppression of genes involved in the BA synthetic pathway (10–12).

SHP, an atypical nuclear receptor that commonly acts as a co-repressor, is considered a key FXR downstream target for many metabolic processes (13–15) in a linear pathway. However, studies with double knockout mice of *Fxr* and *Shp* showed a coordinate role for these receptors in bile acid regulation (16). Examination of *Shp-null* and liver-specific *Shp* knockout (*LShpKO*) mice revealed protective effects against accumulating hepatic lipid levels, albeit through distinct molecular mechanisms (17–19).

Although previous studies to examine the tissue-specific functions of FXR, studies on intestinal SHP using an intestine-specific *Shp* knockout (*IShpKO*) model are lacking. Here we generated, *IShpKO* mice by crossbreeding floxed (*f/f*) *Shp* with mice with a *Cre* recombinase driven by Villin-1 promoter. We examined the impact of intestinal *Shp* deletion on BA homeostasis, which will tease out the contribution of SHP in lipid and fat absorption in the intestine for future studies.

## Materials and Methods

### Generation and genotyping of intestine-specific *Shp* knockout mice

Intestine-specific *Shp* knockout (*IShpKO*) mice were generated by crossing C57/B6 floxed *Shp* (*f/f Shp* (obtained from Dr. Kristina Schoonjas (EPFL, Lausanne Switzerland)) and the B6.Cg-Tg(*Vil1-cre*) 1000Gum/J, obtained from the Jackson Laboratory. *f/f Shp* mice were backcrossed to C57/B6 for ∼10 generations. Mice were genotyped using PCR. Mouse tails were snipped and digested in 100:1 ratio of Direct PCR Lysis Reagent (Tail) (FisherScientific #NC9724951) and Proteinase K (20mg/mL, 10mM Tris-HCl, pH 8.0, 1mM CaCl_2_, and 30% Glycerol) (IBI Scientific # IB05402) overnight 55°C. The enzyme was then inactivated at 95°C for 10 minutes. The digested sample was utilized in GoTaq PCR reactions. The *f/f Shp* and tissue-specific knockout were distinguished based on two primers: Forward primer 5’-TAGTTGCTTGTGGAAAGGACCAACC-3’ and reverse primer 5’-CTAGGAAGTGAAGTGGCCTTGTCTG-3’. These primers produced a product pf 365 base pairs (wild-type), 365/415 base pairs (heterozygous), and 415 base pairs (homozygous). The *Vil1-Cre* transgene was assayed with forward primer 5’-GTGTGGGACAGAGAACAAACC-3’ and reverse primer 5’-ACATCTTCAGGTTCTGCGGG-3’. Cycling parameters were 95°C for 5 minutes, then 35 cycles of 95°C for 30 seconds, 62°C for 1 minute, and 72°C (30 seconds for *f/f Shp*) (1.5 minutes for *Vil1-Cre*), and finally 72°C for 5 minutes. These primers produced a product of 1100 base pairs.

### Animal studies

Mice were maintained, genotyped, and bred in flow cages in a temperature-controlled (23°C) facility on a standard 12-hour light/dark cycles. At 10-12 weeks, *f/f Shp* and *IShpKO* mice were placed on normal chow or 1% cholic acid (CA) (Envigo Teklad, Madison, WI) for either 5 days (acute) or 2 weeks (chronic). Mice were fasted for 6 hours for the acute study, while the mice were maintained in fed-state for the 14-day study prior to sacrifice. The liver, gallbladder, and intestine tissues and serum were collected for gene and protein expression, bile acid assay, and histology. National Institute of Health (NIH) guidelines for use and care of laboratory animals were followed, and all experiments were approved the Institutional Animal Care and Use Committee at the University of Illinois at Urbana-Champaign (Champaign, IL).

### Enterocyte isolation for RNA analysis

The intestine was excised and was cut into thirds (proximal, middle, and distal portions). Each portion was washed with iced cold 1XPBS using a syringe needle (3x 5mL). The intestines were turned inside-out with forceps, cut into 0.5cm pieces, and placed in conical tubes with iced cold 1XPBS and further washed by vigorous shaking for 10-15 seconds, twice. Afterward, the buffer was decanted and iced cold 1XPBS + 2mM EDTA was added. The tubes were laid on a shaker and gently agitated for an 30-60 minutes in 4°C cold room. The enterocytes were dislodged by vigorous shaking for 30 seconds and isolated by filtering through a 70µm strainer (Corning #431751) and pelleted down by centrifugation at 1000g at 4°C. The cell pellet was then washed with iced 1XPBS and pelleted down three times before proceeding to RNA isolation.

### RNA isolation and mRNA quantification

Total RNA from mice livers and the isolated enterocyte from the small intestine were isolated using TRIzol (Invitrogen #15596018) reagent and prepared according to the manufacturer’s instructions. After DNAse treatment (NEBioLabs #M0303S), 2-5µg of RNA was reverse transcribed using random primers (NEBioLabs #S1330S) and Maxima reverse transcriptase kit (ThermoScientific #EP0742). The complementary DNA was diluted to 12.5ng/µL, and 50ng was used for every SYBR green-based qRT-PCR reaction (ThermoScientific #A25778). qRT-PCR primers are provided in Supplemental Table 1.

### Serum blood assay

Retro-orbital bleeding was performed to collect blood from mice in microtainer (FisherScientific #02675186). Blood was coagulated without shaking at room temperature for 30 minutes and then centrifuge at 10,000*g* for 3 minutes to obtain serum. The samples were analyzed according to the Total Bile Acids (NBT Method) test kit (Diazyme #DZ092A-K). 2-4µL of serum was incubated with reagent R1 for 5 minutes at 37°C and then added reagent R2 and incubated for 2 minutes and read at 405nm.

### Tissue bile acid analysis

75-100mg of liver or distal portion of the intestine tissue sample was homogenized in 1mL ice-cold 70% ethanol and incubated for 2 hours in 55-60°C water bath. The samples were then centrifuged at room temperature at 4000rpm. The supernatant was collected and used for bile acid analysis. For biliary bile acid, the intact gallbladder was carefully collected and placed in the eppendorf tube. The gallbladder was then punctured with a needle to release its contents. Tissue and biliary bile acid concentration were analyzed with colorimetric assay as described above.

### Western blot analysis

Protein was extracted from 50-100mg from liver or ileum tissue. Homogenization buffer (7.5mM HEPES pH 7.5, 5mM EDTA, 0.32M sucrose, protease inhibitor (Pierce Protease Inhibitor Mini Tablets, EDTA Free; ThermoFisher Scientific #A32953), and phosphatase inhibitor (Phosphatase Inhibitor Cocktail 3; Sigma Aldrich #P0044) was added to each sample. The samples were homogenized by adding 8-10 1mm beads and bullet blend 5 ⨯ 1 minute with 1 minute on ice in between. Afterward, sodium dodecyl sulfate solution (final conc. was 0.5%) was added into each sample and then sonicated in iced water bath for 15 minutes (60% amplitude, 2-second pulses). Pierce BCA Protein Assay Kit (Thermofisher Scientific #23225) was used to measure the protein concentration. For Western blot, 25-50µg total protein was used. The membrane was incubated with the following antibodies (provided generously by Dr. Paul Dawson’s lab): Rabbit anti-mouse ASBT (1:1000), OSTα (1:1000), and OSTβ (1:1000). Rabbit anti-GAPDH (1:1000; Sigma-Aldrich #G9545) was used as control and goat anti-Rabbit IgG (H+L) secondary antibody (1:5000; ThermoFisher Scientific #31460) was used for all targets.

### Histology

Liver and ileum tissue samples were fixed in 10% neutral-buffered formalin (VWR International #89370-094) and then was embedded in paraffin wax for sectioning (5µm) and processing. Slides were stained with hematoxylin and eosin (Richard-Allan Scientific #7211L and #71311) or alcian blue (Kodak #C14091) and nuclear fast red (Sigma-Aldrich #N8002-5G). Quantification of goblet cells, villi height, and crypt depth were done by averaging 15-18 villi from each sample using ImageJ software. For immunohistochemistry, samples were subjected to antigen retrieval with 10mM sodium citrate pH 6.0 in a microwave for 30 minutes. The samples were cooled at room temperature for 30 minutes, rinsed with water, and quenched with 3% hydrogen peroxide. Slides were incubated with primary antibody rabbit anti-Lysozyme (1:400; Dako, #A0099) and secondary antibody Goat Anti-Mouse (H+L) HRP Conjugate (1:250; BioRad #170-6516). The samples were then incubated with peroxidase (HRP) detection system according to the DAB substrate kit (Vector #SK-4100) and then counterstained with hematoxylin for 15 seconds.

### Tissue clearing and staining reagents

*f/f Shp* and *IShpKO* mice on standard chow were analyzed for this protocol. CUBIC-1 reagent was made by adding 25% (W/V) urea (Sigma #U15-500), 25% (W/V) N, N, N’, N’-tetrakis(2-hydroxypropyl) ethylenediamine (Sigma #122262) and 15% (W/V) Triton X-100 (VWR, #M143) into 0.1M PBS (Lonza #17517Q). CUBIC-2 reagent was made by adding 50% (W/V) sucrose (FisherScientific #BP220-1), 25% (W/V) urea, 10% (W/V) 2,2’,2’-nitrilotriethanol/triethanolamine (Sigma #90279) and 0.1% (V/V) Triton X-100 into 0.1M PBS. Sodium azide stock solution contained NaN_3_ (Sigma #71289) in DI water at a final concentration of 10% (W/V). Blocking solution for tissue staining included 0.1% (V/V) Triton X-100, 4% (V/V) FBS, 0.01% (V/V) (Sigma, #05470), sodium azide and 1:100 rat anti-mouse FcR (rat anti-mouse CD16/32) (Miltenyl Biotec, #130-092-575) to 0,1M PBS. Wash solution contained 0.1% (V/V) Triton X-100 and 0.01% (V/V) sodium azide in 0.1M PBS. Invitrogen^™^ Wheat germ Agglutinin (WGA) conjugated with Alexa Fluor^™^ 594 (ThermoFisher Scientific #W11262) and used at the concentration of 5.0 ug/mL. Invitrogen^™^ DAPI (4’,6-Diamidino-2-Phenylindole, Dihydrochloride) (ThermoFisher Scientific #D1306) and used at the concentration of 1.0 ug/mL in wash solution.

### Tissue clearing

Ileum samples were fixed in 4% paraformaldehyde (PFA) in 0.1M PBS and incubated overnight at 4°C. Samples were rinsed with sterile 0.1M PBS three times with rocking at room temperature for 15 minutes each to remove PFA. Tissue samples were immersed in CUBIC-1 reagent at 37°C with gentle shaking and daily buffer exchanges until the samples were completely decolorized (17 days for the current study). Samples were washed three times with 0.1M PBS for 30 min each at room temperature with gentle shaking. Tissue samples were then immersed in CUBIC-2 reagent for 21 days at 37°C with gentle rocking until transparent. Samples were stored in CUBIC-2 with 0.01% (V/V) sodium azide in the dark until ready for immunostaining. Tissues were imaged before and after clearing in order to record the levels of tissue transparency.

### Tissue staining and confocal microscopy

The samples were washed three times with 0.1M PBS for 30 minutes each at room temperature with gentle shaking and then blocked overnight at 4°C with gentle agitation. The samples were stained with 5.0 ug/mL WGA conjugated with Alexa Fluor^™^ 594 (centrifuge at 5,000 rpm for five minutes before use to minimize protein aggregation) in blocking solution (minus anti-mouse FcR) for 3 days at room temperature with rotation. WGA stained samples were washed five times with wash solution at room temperature with rotation over 5 hours. Nuclei were stained with DAPI by adding 1.0 ug/mL DAPI stain solution and incubated overnight at 4°C with agitation. Samples were washed with washing solution three times at room temperature with rotation for 30 minutes in total, then immersed in CUBIC-2 reagent for 24 hours at room temperature before mounting. Samples were mounted between two No.1 coverslips (Electron Microscopy Sciences #63768-01) separated by adhesive/adhesive silicone isolators (Electron Microscopy Sciences #70336-70) in CUBIC-2 solution and imaged via a Zeiss LSM-700 confocal microscope.

### Statistical analyses

Data are expressed as ± SD. For comparisons between two groups, student’s unpaired two tail *t*-test was used. For comparisons for more than two groups, two-way ANOVA was performed, followed by the Bonferroni *post hoc* test. Statistical significance was determined as *p* < 0.05. Each statistical significance difference (*p<0*.*05)* are indicated by bars with their calculated *p-*value determined by GraphPad Prism. Outliers were determined by Grubb’s test and removed from the analysis.

## Results

### Ileal *Shp* expression controls basal and BA-induced alterations in *Fgf15* levels and BA levels

Region-specific expression of *Shp* and other essential BA regulators and transporters were mapped in the small and large intestines of mice. qPCR analysis revealed that the expression of *Shp, Fxrα*, fibroblast growth factor 15 (*Fgf15*), apical sodium-dependent bile acid transporter (*Asbt*), *and* organic solute transporter *α/β* (*Ostα/β*) was highly localized in the distal portion of the small intestine compared to the other regions (Suppl. Fig. 1A-F) (14,20). The large intestine also expressed *Shp, Fxrα*, and *Asbt* but not to the same extent as the ileum. Based on the expression patterns, we focused the rest of the study on the distal portion of the small intestine.

**Figure 1.**
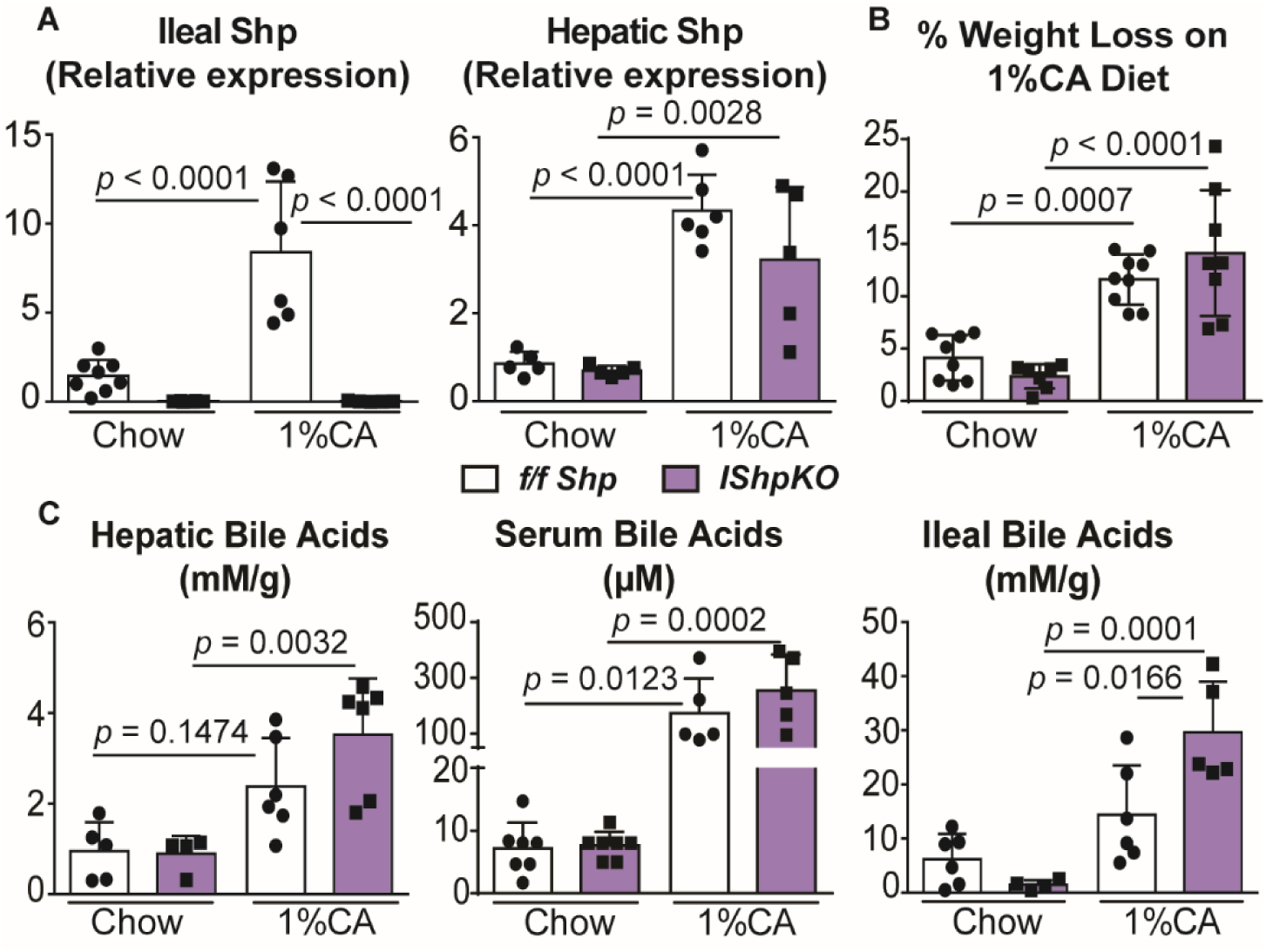
Deletion of intestinal *Shp* altered bile acid concentration when challenged with a 1%CA diet. (A) Relative mRNA expression of ileal and hepatic *Shp* confirmed that the deletion was restricted to the intestine. (B) Reduction in body weight was similar between *f/f Shp* and *IShpKO* after 5-day 1%CA diet in 10- to −12-week-old mice. (C) Quantitative analysis of total bile acid concentration in the liver, serum, and ileum. Data are represented as mean ±SD, n=6-8, and analyzed with two-way ANOVA, Bonferroni’s test.

We generated homozygous *IShpKO* mice and confirmed the deletion of *Shp* was restricted to the intestine by genotyping and qPCR analyses (Suppl. Fig. 2A and Fig. 1A). *Shp* was not detected in the intestine, while hepatic *Shp* transcript was expressed normally with no compensatory changes. *IShpKO* intestines also maintained similar localization to control mice, indicating that the region-specific distribution of BA regulators and transporters is independent of SHP. When challenged with a chronic 1%CA diet, the loss of body weight in *IShpKO* mice rebounded after the 5^th^ day, with no significant difference from the *f/f Shp* throughout the remainder of the 14-days (Suppl. Fig. 2B and Fig. 1B). To further evaluate significant weight loss on day 5, we investigated the response of *IShpKO* mice during acute (5 days) and chronic (14 days) BA diet treatment. As expected, the acute CA diet increased hepatic and serum BAs in both *f/f Shp* mice and *IShpKO* mice. Although lower basal concentrations of ileal BA concentration were observed, *IShpKO* mice accumulated more ileal BA upon 5 days of CA diet (∼20 fold) compared to *f/f Shp* mice (∼2 fold) (Fig.1C). Conversely, a chronic CA diet resulted in comparable hepatic and ileal BAs between the two genotypes, but higher serum BA concentrations were seen in *IShpKO* mice (Fig. 1C and Suppl. Fig. 2C).

**Figure 2.**
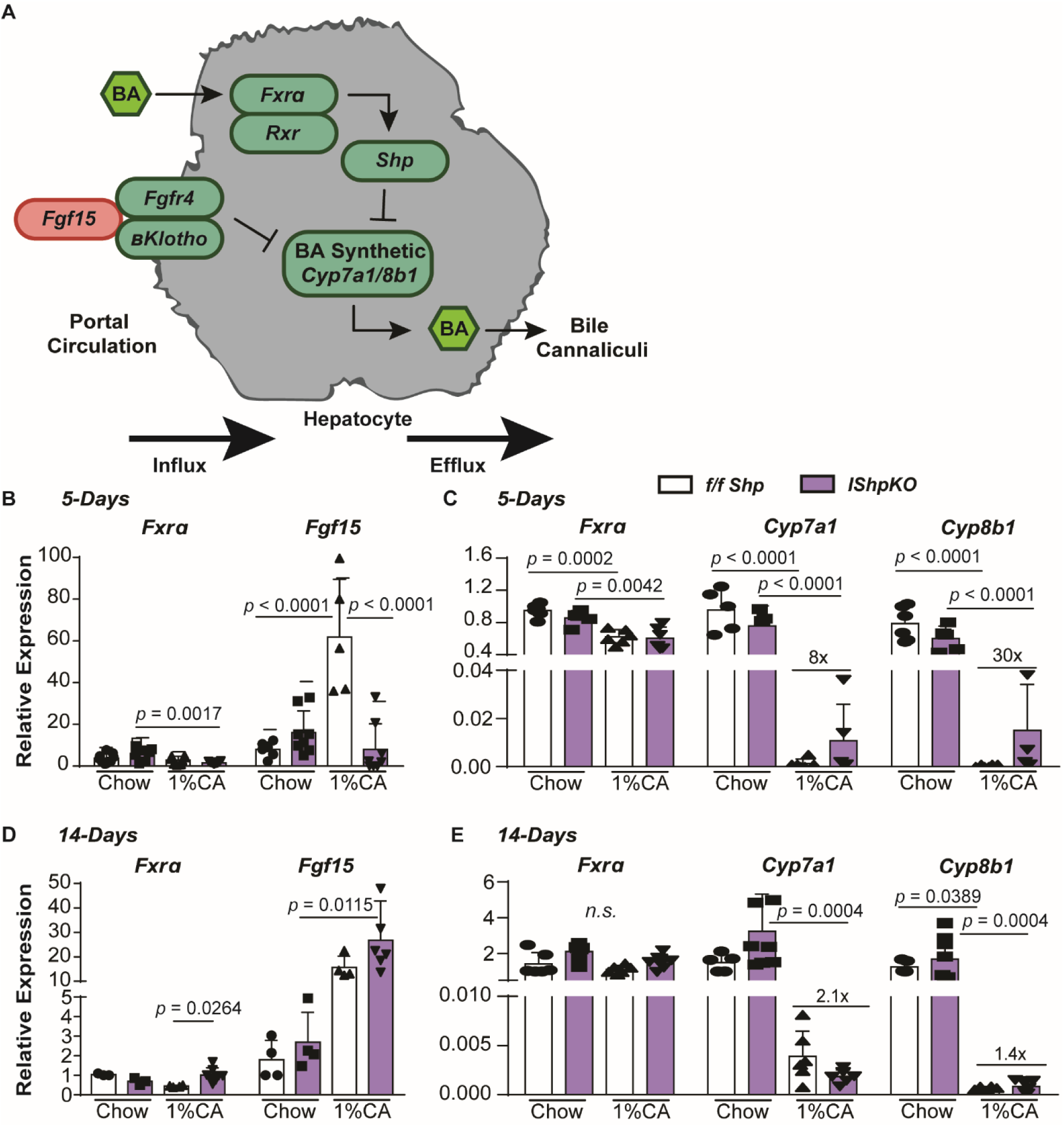
The ileal FGF15 signaling response to the CA diet is defective in *IShpKO* mice. f/f Shp and *IShpKO* mice were challenged with an acute (5 days) and chronic (2 weeks) 1%CA or standard chow diet. (A) Schematic mechanism of liver bile acid regulation. (B) Acute CA diet challenge with 6-hour fasting decreased *Fgf15* mRNA expression. (C) Suppression of hepatic bile acid synthetic genes, *Cyp7a1* and *Cyp8b1*, upon CA diet was reduced in *IShpKO* mice compared to *f/f Shp* control. (D) In contrast, the chronic CA diet increased *Fgf15* mRNA expression. (E) *IShpKO* mice showed a lesser ability to suppress bile acid synthetic gene expression in response to the chronic CA diet. Data are represented as mean ±SD, *n*=6-8 and analyzed with two-way ANOVA, Bonferroni’s test.

### Absence of intestinal SHP leads to defects in the BA enterohepatic recirculation machinery

*IShpKO* mice displayed a higher liver-to-body weight ratio both under normal and CA diet-fed conditions but did not show obvious histological changes in the liver (Suppl. Fig. 3A-C). Acute CA challenge caused significant increases in ALT and AST levels (Suppl. Fig. 3D), markers for liver injury, which correlates with the higher serum BA concentration in both *f/f Shp* and *IShpKO* mice (Fig. 1C and Suppl. Fig. 2C). Acute CA diet led to significantly higher levels of AST in *IShpKO* compared to *f/f Shp* mice.

**Figure 3.**
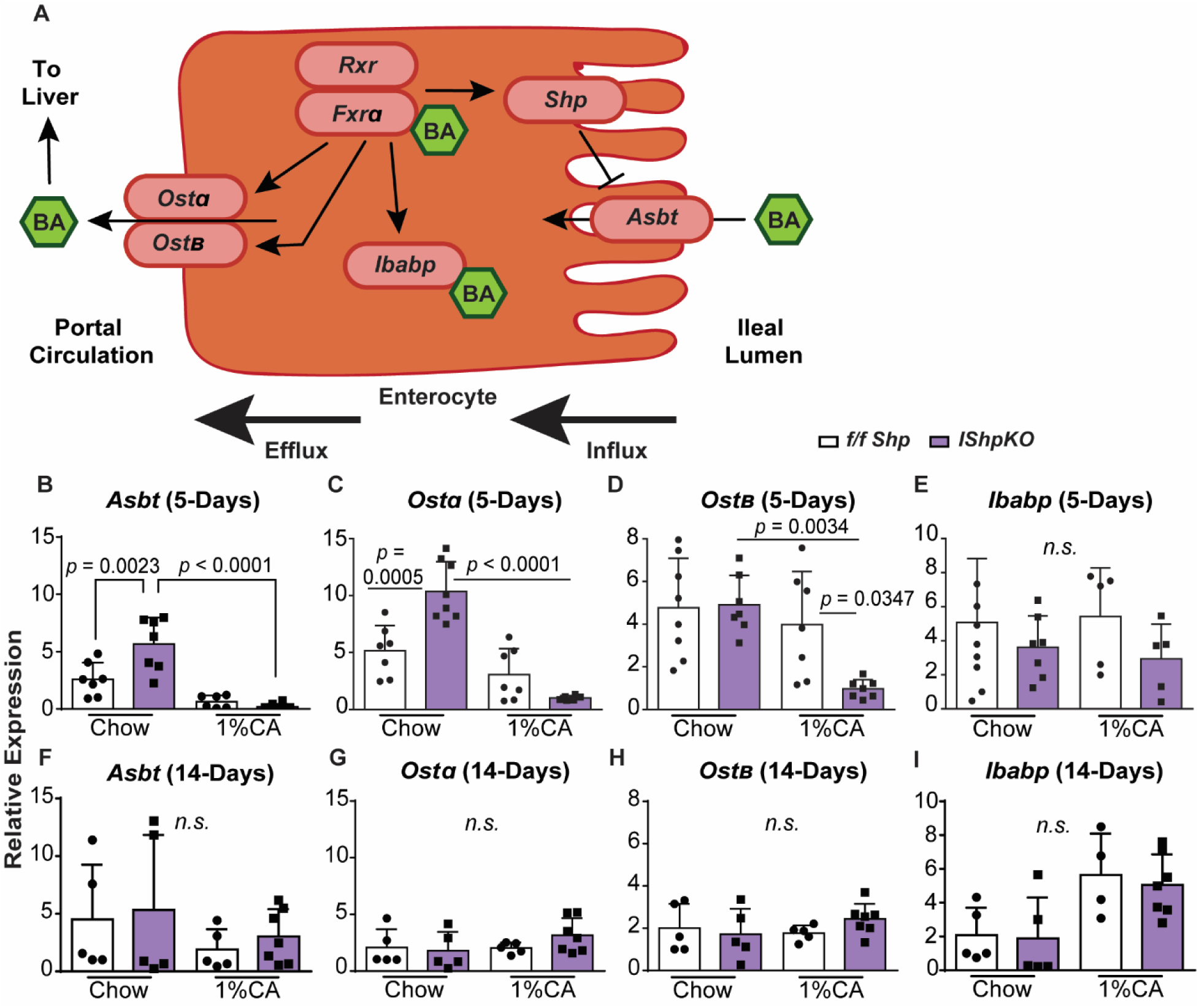
Adaptive regulation of ileal bile acid transporters is observed in the chronic CA diet when intestinal *Shp* expression is deleted. mRNA expression of bile acid transporters in the intestine in *IShpKO* compared to *f/f Shp* mice under chow and CA-fed conditions. (A) Schematic mechanism of ileal bile acid regulation. (B-E) 5-day CA diet and 6 hours fast in *IShpKO* mice showed ileal bile acid transporter gene suppression. (F-I) 14-day CA and kept in a fed state. Data are represented as mean ±SD, *n*=5-8 and analyzed with two-way ANOVA, Bonferroni’s test.

FXR and SHP orchestrate the crosstalk between the intestine and the liver via the *Fgfr4-Fgf15* axis to control BA homeostasis (Fig. 2A) (3,7) (3,14,21). Briefly, FXRα induces FGF15 expression in the terminal ileum. FGF15 circulates and binds to FGFR4 and β-KLOTHO in the liver and, in coordination with SHP, represses BA synthesis (22– 25). So, we investigated this intestine-liver communication when intestinal *Shp* is deleted and found intact basal expression of *Fxr* and *Fgf15* in the intestine (Fig. 2B and 2D). Upon CA diet, *Fxr* expression was unaltered, whereas *Fgf15* was induced in *f/f Shp* mice, while *Fxr* expression was reduced and *Fgf15* induction was completely lost in *IShpKO* mice (Fig. 2B). This intriguing result led us to examine the hepatic expression of *Fxr* and BA synthesis genes in these contexts. *Floxed Shp* control animals showed the expected negative feedback response to BA challenge with reduced expression of BA synthesis genes-*Cyp7A1* and *Cyp8b1*. In contrast, *IShpKO* mice showed blunted suppression of BA synthesis genes, consistent with reduced ileal *Fgf15* expression.

We were surprised to find that the defects in the ileal *Fxr-Fgf15* expression were normalized with long-term CA feeding in *IShpKO* mice (Fig. 2D). In line with this finding, the suppression of *Cyp7A1* and *Cyp8b1* was also not blunted to the same extent as short-term feeding in the *IShpKO* livers (Fig. 2C and 2E). Thus, we find two distinct expression patterns of *Fgf15* depending on the length of CA diet in *IShpKO* ileum. One caveat is that the mice in the 5-day experiment were analyzed after 6 hours of fast, whereas the 14-day CA diet experiment was analyzed without fasting. Therefore, we repeated the 5-day CA-diet experiment and examined *Fgf15* gene expression in the fed-state. Surprisingly, we found no difference in the induction of *Fgf15* gene expression in *f/f Shp* and *IShpKO* mice, indicating the loss of *Fgf15* expression in *IShpKO* is subsequent to short-term CA diet and 6-hour fast (Suppl. Fig. 4A-G). Further examination revealed that, fasting results in overall elevated expression of *Fgf15* than the fed-state despite the fold induction in response to CA-diet being comparable. These data indicate that the loss of intestinal SHP in response to CA-mediated increase in *Fgf15* expression post-6 hour fasted state.

**Figure 4.**
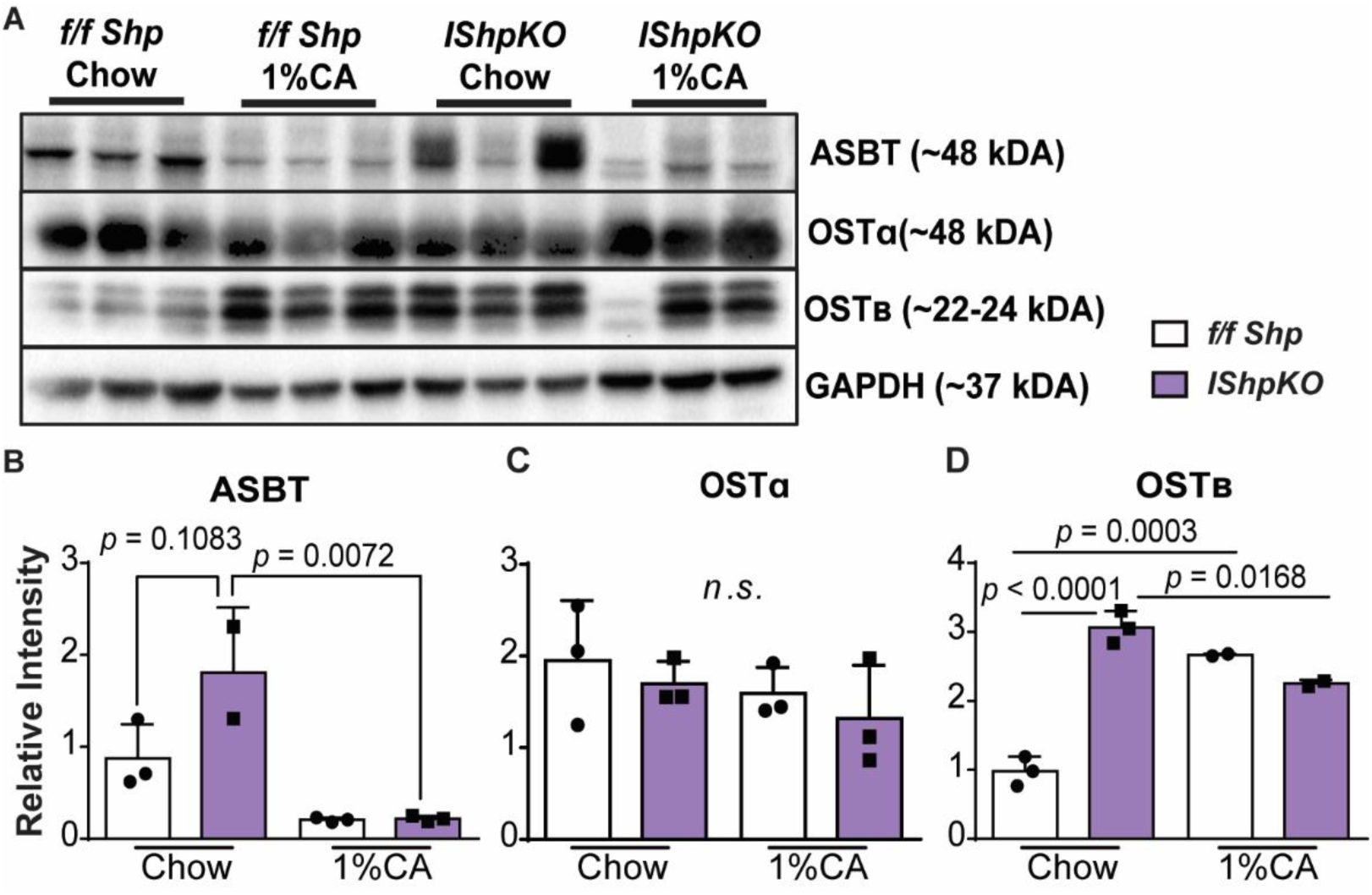
*IShpKO* mice exhibit different levels of Ileal bile acid transport proteins. (A) Western blot of ASBT, OSTα, and OSTβ showed increased expression of some of the transporter protein expression in the absence of ileal SHP. (B-D) Quantification of the protein intensity level of ASBT, OSTα, and OSTβ. Data are represented as mean ±SD, *n*=3. Two-way ANOVA, Bonferroni’s test.

### Ileal BA transporters are misregulated in the absence of intestinal SHP

The terminal ileum is essential for the reabsorption of BAs. ASBT (apical sodium-dependent bile acid transporter) is critical for the uptake of BA from the intestinal lumen and OSTα/β (organic solute transporter) heterodimer facilitate BA export to portal circulation (26–28). SHP has been shown to regulate ASBT negatively and OSTα/β through the FXR-SHP-LRH-1(liver receptor homolog-1) axis, revealing its critical role in controlling BA concentration in ileal cells (27,29–31) (Fig. 3A). Therefore, we assessed the expression of these ileal BA transporters and we found that the *Asbt* and *Ostα* transcripts were higher in *IShpKO* ileum, but the expression of *Ostβ* and ileal bile acid-binding protein (*Ibabp*) were comparable to that of the *f/f Shp* mice (Figure 3B-E). As expected, the acute CA diet decreased the expression of *Asbt* and *Ostα/β* in *f/f Shp* mice. Surprisingly, this suppression was maintained in the *IShpKO* mice. However, *Ibabp* levels were comparable and remained unaltered between the *IShpKO* and *f/f Shp* mice after the CA diet. Intriguingly, the expression of BA transporters in fed-state mice was maintained regardless of diet except for *Ibabp*, which increased in both groups during chronic CA diet (Fig. 3F-I). These data suggest that the BA-enterohepatic recirculation machinery may be misregulated initially during BA excess but then adapts during the chronic CA diet.

Consistently with the qPCR data, western blot analyses revealed that ileal ASBT protein expression was higher in *IShpKO* mice than in *f/f Shp* mice even under chow-fed conditions. Despite the loss of SHP, acute CA-mediated suppression of ASBT protein levels was maintained (Fig. 4A-B). Although *Ostα* increased transcript levels, OSTα protein levels were not different between the two groups of mice under chow and CA-fed conditions (Fig. 4C). On the other hand, OSTβ protein expression in the ileum was induced in the *IShpKO* group and this increase was to the same extent to that of CA-fed *f/f Shp* mice. Further, OSTβ protein expression remained high when intestinal SHP was absent regardless of diet (Fig. 4D). *Ibabp*, a protein responsible for trafficking BA from the apical to the basolateral side of the enterocyte, was found to be expressed at similar levels between all the groups during the acute CA diet (Fig. 3E). In contrast, the chronic CA challenge increased the expression of *Ibabp* transcript (Fig. 3I). Because the expression of CA transporters was changed in the ileum, we examined their hepatic expression. Since hepatic *Shp* expression was intact, hepatic transporters retained their regulation (Suppl. Fig. 5A-G) except for Na^+^-taurocholate cotransporting polypeptide (*Ntcp*) gene expression, which was higher in chronic CA-fed *IShpKO* mice (Suppl. Fig. 5B).

**Figure 5.**
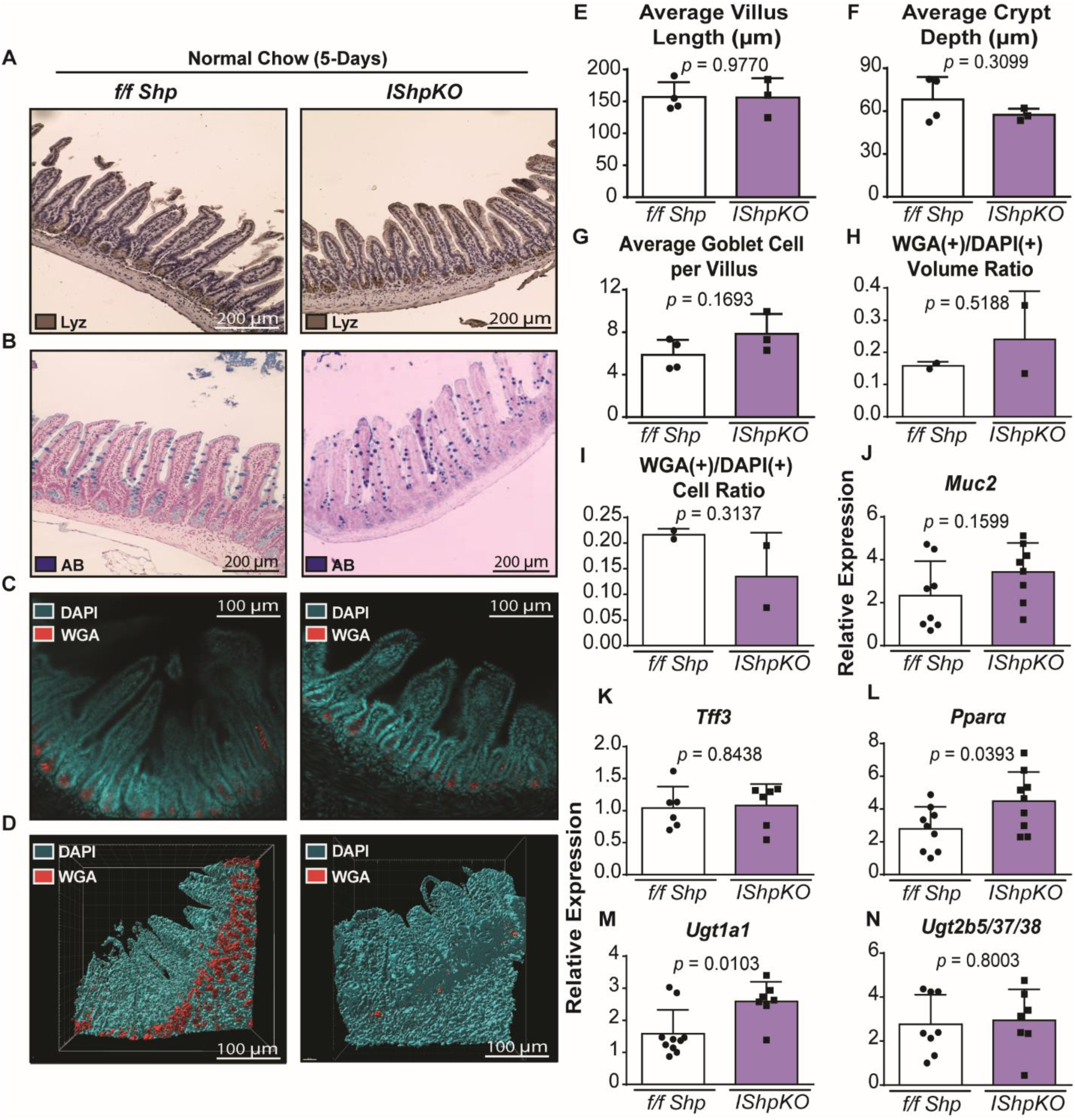
Characterization of Intestinal morphology in *IShpKO* mice. Small intestines of the adult *f/f Shp* control and *IShpKO* mice were analyzed on chow diet. (A) Paneth cell population was examined using lysozyme staining, which showed no difference in staining intensity. (B) Goblet cells were stained using alcian blue. (C) Z-slice of fluorescence imaging and (D) surface model representation of tissue cleared ileum after stained with WGA and DAPI. Quantification of (E) average villus length, (F) average crypt depth, and (G) average goblet cell count per villus in *IShpKO* under chow diet. (H) WGA(+) cells-to-DAPI(+) cells ratio and (I) WGA(+)-to-DAPI(+) volume ratio was calculated by Imaris software, *n*=3-4. (See also supplemental movies 1, 2, 3, and 4). Goblet cell markers (J) *Muc2* and (K) *Tff3* were not significantly different from *f/f Shp* and *IShpKO*. (L-N) *Pparα* and *Ugt* mRNA transcript levels were increased when intestinal *Shp* is deleted, *n*=6-8. Data are represented as mean ±SD, Unpaired *t*-test.

### *Pparα-Ugt* axis is upregulated during chronic CA challenge

Next, we examined the intestinal epithelial structure and mucin levels as they are altered when there is an accumulation of BAs (32). Notably, SHP has been recently identified as a goblet cell marker (33) and can regulate immune function and cell proliferation (34,35) and was recently identified as. Under normal chow-fed conditions, SHP loss in the intestine is tolerated with no change in Paneth cells or goblet cells (Fig. 5A-D) and does not alter the average villus length (Fig. 5E) and crypt depth (Fig. 5F). Using tissue clearing and WGA staining (Suppl. Fig. 6), we analyzed the 3D distribution of goblet cells within intact regions of the ileum and observed no significant difference in the WGA(+)-to-DAPI(+) volume (Fig. 5G) and cell ratio (Fig. H) (See also movies 1, 2, 3 and 4) under basal conditions. This correlated well with no changes in the expression of goblet cell markers, mucin 2 (*Muc 2*) (Fig. 5J) and trefoil factor 3 (*Tff3*) (Fig. 5K) in the *IShpKO* mice. We also investigated the expression of peroxisome proliferator-activated receptors α (*Pparα*) and UDP-glucuronosyltransferase (*Ugt*) involved in metabolic BA elimination (36) in the ileum (Fig. 5L-N) and found increased expression in *IShpKO* compared to chow fed *f/f Shp*. However, the acute CA-diet led to shortened and clubbed villi resulting in reduced lengths, crypts with varying widths, and possible injury (Fig. 6A-B) in *IShpKO* mice. We also observed a reduction in goblet cell counts (Fig. 6E) along with reduced gene expression of *Tff3* but not *Muc2* (Fig. 6F-G). *Pparα, Ugt1a1, Ugt2b/37/38* were unaltered in *IShpKO* mice compared to *f/f Shp* control (Fig. 6H-J).

**Figure 6.**
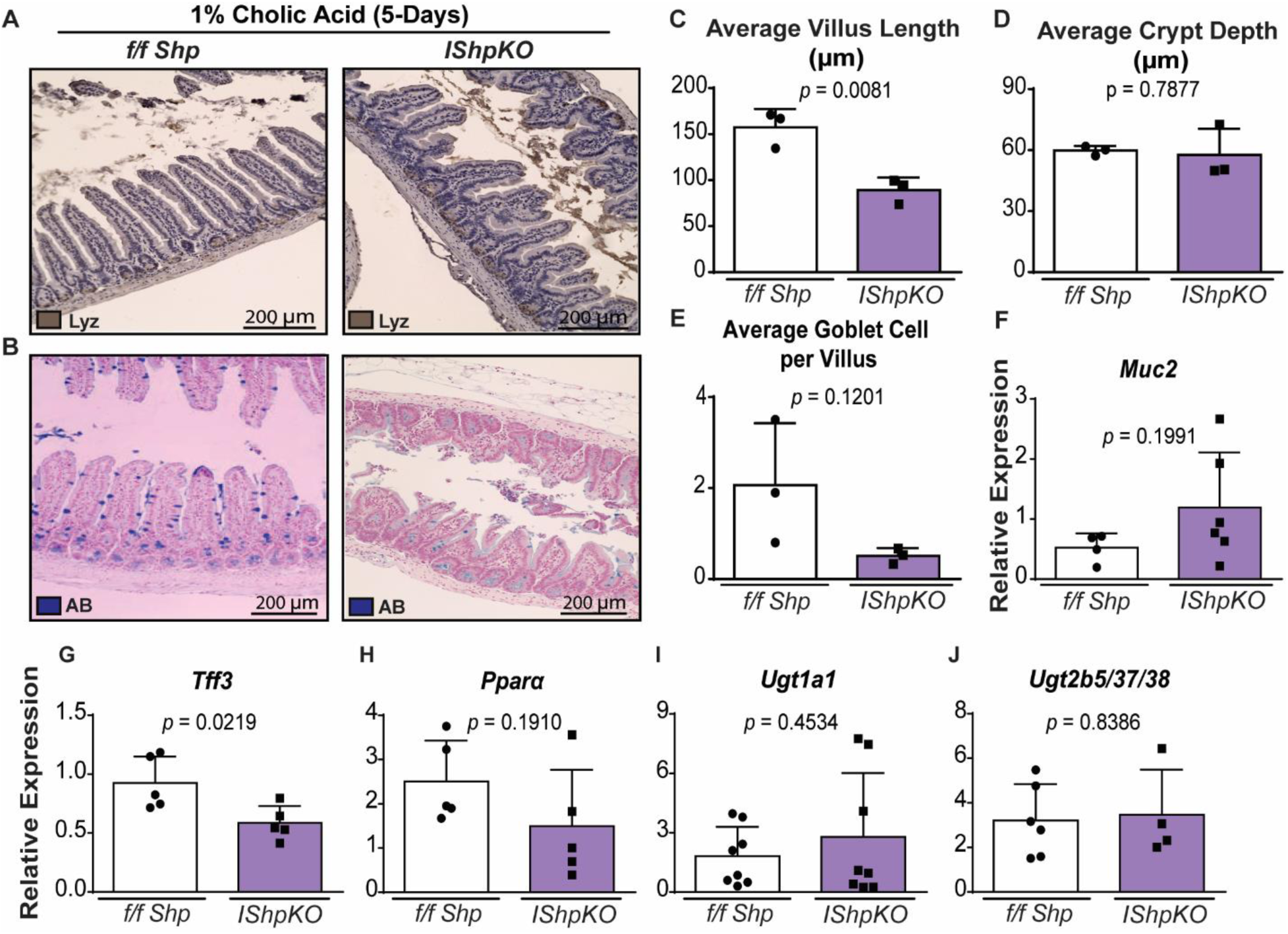
Alterations in intestinal morphology and *Pparα*-*Ugt* axis when mice are challenged with acute CA diet. *f/f Shp* and *IShpKO* mice were challenged with either chow or acute 1%CA enriched diet. (A) No difference in Paneth cell count was detected with lysozyme staining between control and *IShpKO mice*. (B) Histomorphological analysis using alcian blue staining showed lower goblet cell counts and smaller villi in *IShpKO* mice upon CA challenge. Quantification of (C) average villus length, (D) average crypt depth, and (E) average goblet cell count per villus in *IShpKO*. (F) *Muc2* gene expression was higher than the *f/f Shp* control. (G) *Tff3* mRNA transcript level was significantly higher in *IShpKO* but (H) *Pparα* mRNA levels were decreased. (I-J) The expression of *Ugt* genes regulating glucuronidation that can help detoxify bile acids were not altered in the ileum. Data are represented as mean ±SD, *n*=3-6. Unpaired t-test.

Next, we investigated whether chronic CA challenge alters intestinal morphology in *IShp*KO. Ileal intestinal morphology showed no sign of abnormality or injury (Suppl. Fig. 7A-B) in contrast to acute CA diet (Fig. 5B and 6B). Similarly, the villi length (Suppl. Fig. 7C), crypt depth (Suppl. Fig. 7D), and goblet cell number (Suppl. Fig. 7E) were unaffected in *IShpKO* mice. This result was contrary to our expectation that the chronic CA diet would have a worsen the acute CA-mediated defects. Instead, *IShpKO* mice rebound and display a comparable response to that of the *f/f Shp* control mice. To investigate a compensatory mechanism, the *Pparα-Ugt* axis transcript levels were examined and found to be induced (Suppl. Fig. 7E-I). These results suggest that the *IShpKO* mice may handle long-term exposure to excess BA by increasing *Pparα* and *Ugt* expression.

## Discussion

Despite essential roles for SHP in controlling the enterohepatic BA recirculation and homeostasis, intestine-specific studies are lacking. This is the first *IShpKO* study, wherein we report a role for *Shp* in regulating goblet cells in addition to biliary homeostasis. As previously reported, we found that many essential BA regulators and transporters are localized in the ileal portion of the small intestine (1,37). Ileal analysis of the *f/f Shp* and *IShpKO* under the chow revealed a significant increase in *Asbt and Ostα* at the mRNA level and ASBT and OSTβ at the protein level in *IShpKO* mice. This is consistent with SHP being a corepressor. Previous studies focusing on the heterodimerization, trafficking, and function of OSTα/β (31,38) revealed the β-subunit of OSTβ is essential for dimerizing with OSTα, trafficking to the plasma membrane, and exporting BA to portal circulation (39). The heterogeneous expression of OSTα/β at the mRNA and protein level when intestinal SHP is absent highlights a regulatory function of SHP at basal and abnormally high BA conditions.

Contrary to proposed mechanisms that SHP suppresses ASBT levels upon excess BAs (29,40), we found this suppression intact in *IShpKO* mice. Further, mechanistic studies have shown that LRH-1 is not essential for BA-mediated ASBT inhibition (41), countering the notion that the FXR-SHP-LRH-1 signaling axis is required of this process. We speculate that pathways (15,42) mediated by either PPARα, glucocorticoid receptor (43), or hepatocyte nuclear factor 1 homeobox A (HNF-1α) (44,45) may compensate for the absence of SHP.

During the acute CA diet, mice were put on a 6-hour fast before sacrificing to allow for gallbladder filling and intestinal emptying of feces, unlike the chronic CA diet experiment where the animals remained in a fed-state. Fasting has been known to regulate BA biosynthesis where *Cyp7a1* was significantly induced at the mRNA level in mice after 24 hours of fasting (46,47). This was also observed in rats, where *Cyp7a1* levels remained constant and then significantly increased after 14 hours of fasting (48). This variable resulted in an unexpected yet exciting observation as fasting has been shown to produce a broad range of changes (49). It was previously shown in broiler chickens that fasting for 4-12 hours can modify the intestinal function and cause alterations to the intestinal morphology and goblet cell number (50,51). Although 6 hours of fasting in *IShpKO* mice did not alter villi length or goblet cell numbers under normal chow conditions, CA-feeding resulted in changes to both these parameters when SHP was deleted in the intestine.

It would be interesting to evaluate whether fasting acts, as an additional stimulus and exacerbates the *IShpKO* mice chronic CA-diet phenotype. SHP expression in the intestine is necessary to maintain normal expression of BA transporters such that loss of SHP leads to increased BA transporters expression. It is possible that the increase in intestinal BAs may result in higher expression of the transporters to reduce BA burden. While ASBT was upregulated in the absence of intestinal SHP under basal conditions, acute CA-mediated suppression of ASBT was shown to be SHP-independent, suggesting an alternative pathway to reduce BA reabsorption into enterocytes. Similarly, OSTβ was upregulated in the absence of intestinal SHP under chow diet, but the protein level for OSTβ was maintained during acute CA excess. Therefore, we posit since OSTβ protein expression was already higher in *IShpKO* mice than *f/f Shp*, it does not get induced further when challenged with CA-diet. These results suggest that intestinal SHP plays an essential role in regulating intestinal architecture, BA uptake and transport (32), goblet cell function, and overall intestinal function/homeostasis (35,52).

## Acknowledgments

Dr. Kristina Schoonajans provided the floxed SHP mice at the Ecolé Polytechnique Féderalé de Lausanne, Lausanne, Switzerland. We thank Dr. Paul Dawson at the Emory University School of Medicine for initial discussions and generously sharing the antibodies that recognize ASBT, OSTα, and OSTβ antibodies. Ms. Jannette Rodriguez-Otero helped Ryan Riessen with histology and analysis in the early stages.

## Author contributions

J.T.N., R.R., and T.Z. did the experiments and data analysis; J.T.N, R.R., and T.Z., C.K., and S.A. did the interpretation of data, and J.T.N. and S.A. wrote the manuscript; J.T.N., T.Z., C.K., and S.A. helped with editing the manuscript. S.A. was responsible for acquiring the funds, study design, and study supervision.

## Figure Legends

**Supplemental Figure 1. Intestinal bile acid regulators and transporters are localized in the distal portion of the intestine**. The small intestine was cut into thirds (proximal (near the stomach), middle, and distal (near the cecum) for determining the localization of bile acid transporter and regulators. At basal conditions, there was no change in mRNA transcript levels between the *f/f Shp* and *IShpKO* mice on standard chow diet. *Shp* was not detected throughout the intestine. All samples collected between 12:00pm – 2:00pm. Data are represented as mean ±SD, *n*=5. Two-way ANOVA, Bonferroni’s test.

**Supplementary Figure 2. Adaptive bile acid regulation is observed in *IShpKO* fed chronic CA diet**. (A) Genotyping PCR confirms the presence of both floxed allele and the *Villin1-Cre* allele in *IShpKO* mice. (B) Weight change was monitored over the course of 14-day CA diet. (C) Bile acid concentrations were quantified from the liver, serum, and ileum. Data are represented as mean ±SD, *n*=5-12. Two-way ANOVA, Bonferroni’s test.

**Supplementary Figure 3. *IShpKO* mice exhibited normal liver architecture**. Histomorphological analysis of both f/f Shp and *IShpKO* under (A) chow and (B) 14-day CA diet showed healthy liver architecture with no obvious injury. (C) The liver-to-body weight ratio was induced as expected by the introduction of chronic CA diet. (D) ALT and

AST levels were measured after 5-day CA diet. Data are represented as mean ±SD, *n*=6-12 and analyzed with a two-way ANOVA, Bonferroni’s test.

**Supplementary Figure 4. Intestinal SHP has no effect on bile acid transporters in mice after 5-day CA diet ad-lib state**. (A-G) mRNA from enterocytes of both *f/f Shp* and *IShpKO* was analyzed with qPCR. Gene expression of transporters regulators involved in BA circulation and homeostasis were not changed between control and *IShpKO* mice. Data are represented as mean ±SD, *n*=7-11 and analyzed with a two-way ANOVA, Bonferroni’s test.

**Supplementary Figure 5. Liver bile acid transporters are unaltered in *IShpKO* mice**. mRNA from liver tissue of both control *f/f Shp* and *IShpKO* mice was analyzed with qPCR. Gene expression of transporters involved in BA import and export were not changed between control and *IShpKO* mice. (A) Schematic mechanism of bile acid transportation. (B-C) Acute and chronic CA diet comparison between *Ntcp and Bsep*. (E-G) Hepatic BA transporters after chronic CA diet. Data are represented as mean ±SD, *n*=6-12 and analyzed with a two-way ANOVA, Bonferroni’s test.

**Supplementary Figure 6. Method for tissue clearing using CUBIC**. (A) Flow chart showing the process for tissue clearing and staining using CUBIC. (B) Before and after images of intestinal tissue using CUBIC.

**Supplementary Figure 7. Intestinal goblet cell counts were maintained during chronic CA diet**. (A-E) Histomorphological analysis using alcian blue staining revealed no statistical difference in the number of goblet cells and crypt depth but there was increase in villi length in *IShpKO* mice after chronic CA diet. Data are represented as mean ±SD, *n*=3-5. Unpaired t-test. (F-J) The expression of *Pparα* and *Ugt* genes regulating glucuronidation of metabolic bile acid for elimination were upregulated in the ileum. Data are represented as mean ±SD, *n*=3-6 and analyzed with a two-way ANOVA, Bonferroni’s test.

## References

1. Dawson PA, Karpen SJ. Intestinal transport and metabolism of bile acids. J. Lipid Res. 2014;56(6):1085–1099.

2. Schonewille M, De Boer JF, Groen AK. Bile salts in control of lipid metabolism. Curr. Opin. Lipidol. 2016;27(3):295–301.

3. Russell DW. Fifty years of advances in bile acid synthesis and metabolism. J. Lipid Res. 2009;50(Supplement):S120–S125.

4. Taoka H, Yokoyama Y, Morimoto K, Kitamura N, Tanigaki T, Takashina Y, Tsubota K, Watanabe M. Role of bile acids in the regulation of the metabolic pathways. World J. Diabetes 2016;7(13):260.

5. Shapiro H, Kolodziejczyk AA, Halstuch D, Elinav E. Bile acids in glucose metabolism in health and disease. J. Exp. Med. 2018;215(2):383–396.

6. Tiratterra E, Franco P, Porru E, Katsanos KH, Dimitrios K. Role of bile acids in inflammatory bowel disease. Ann. Gastroenterol. 2018;31:266–272.

7. Li, Tiangang, Chiang J. Nuclear receptors in bile acid metabolism. Drug MEtab Rev 2013;45(1):145–155.

8. Hylemon PB, Zhou H, Pandak WM, Ren S, Gil G, Dent P. tBile acids as regulatory molecules. J. Lipid Res. 2009;50(8):1509–1520.

9. Sinal CJ, Tohkin M, Miyata M, Ward JM, Lambert G, Gonzalez FJ. Targeted disruption of the nuclear receptor FXR/BAR impairs bile acid and lipid homeostasis. Cell 2000;102(6):731–744.

10. Schmitt J, Kong B, Stieger B, Tschopp O, Schultze SM, Rau M, Weber A, Müllhaupt B, Guo GL, Geier A. Protective effects of farnesoid X receptor (FXR) on hepatic lipid accumulation are mediated by hepatic FXR and independent of intestinal FGF15 signal. Liver Int. 2015;35(4):1133–1144.

11. Kim I, Ahn S-H, Inagaki T, Choi M, Ito S, Guo GL, Kliewer SA, Gonzalez FJ. Differential regulation of bile acid homeostasis by the farnesoid X receptor in liver and intestine. J. Lipid Res. 2007;48(12):2664–2672.

12. Li T, Matozel M, Boehme S, Kong B, Nilsson LM, Guo G, Ellis E, Chiang JYL. Overexpression of cholesterol 7α-hydroxylase promotes hepatic bile acid synthesis and secretion and maintains cholesterol homeostasis. Hepatology 2011;53(3):996–1006.

13. Båvner A, Sanyal S, Gustafsson Jå, Treuter E. Transcriptional corepression by SHP: Molecular mechanisms and physiological consequences. Trends Endocrinol. Metab. 2005;16(10):478–488.

14. Lu TT, Makishima M, Repa JJ, Schoonjans K, Kerr TA, Auwerx J, Mangelsdorf DJ. Molecular basis for feedback regulation of bile acid synthesis by nuclear receptos. Mol. Cell 2000;6(3):507–515.

15. Wang L, Lee Y-K, Bundman D, Han Y, Thevananther S, Kim C-S, Chua SS, Wei P, Heyman RA, Karin M. Redundant pathways for negative feedback regulation of bile acid production distinct oxysterol 7-hydroxylase (CYP7B) (Axelson et al. Dev. Cell 2002;2:721–731.

16. Anakk S, Watanabe M, Ochsner SA, Mckenna NJ, Finegold MJ, Moore DD. Combined deletion of of Fxr and Shp in mice induces Cyp7a1 and results in juvenile onset cholestasis. J Clin Invest 2011;121(1):86–95.

17. Hartman HB, Lai K, Evans MJ. Loss of small heterodimer partner expression in the liver protects against dyslipidemia. J. Lipid Res. 2008;50(2):193–203.

18. Akinrotimi O, Riessen R, Vanduyne P, Park JE, Lee YK, Wong L, Zavacki AM, Schoonjans K, Anakk S. Small heterodimer partner deletion prevents hepatic steatosis and when combined with farnesoid X receptor loss protectes agasinst type 2 diabetes in mice. Hepatology 2017;66(6):1854–1865.

19. Magee N, Zou A, Ghosh P, Ahamed F, Delker D, Zhang Y. Disruption of hepatic small heterodimer partner induces dissociation of steatosis and inflammation in experimental nonalcoholic steatohepatitis. J. Biol. Chem. 2020;295(4):994–1008.

20. Goodwin B, Jones SA, Price RR, Watson MA, McKee DD, Moore LB, Galardi C, Wilson JG, Lewis MC, Roth ME, Maloney PR, Willson TM, Kliewer SA. A regulatory cascade of the nuclear receptors FXR, SHP-1, and LRH-1 represses bile acid biosynthesis. Mol. Cell 2000;6(3):517–526.

21. Kuipers F, Bloks VW, Groen AK. Beyond intestinal soap - Bile acids in metabolic control. Nat. Rev. Endocrinol. 2014;10(8):488–498.

22. Inagaki T, Choi M, Moschetta A, Peng L, Cummins CL, McDonald JG, Luo G, Jones SA, Goodwin B, Richardson JA, Gerard RD, Repa JJ, Mangelsdorf DJ, Kliewer SA. Fibroblast growth factor 15 functions as an enterohepatic signal to regulate bile acid homeostasis. Cell Metab. 2005;2(4):217–225.

23. Kim YC, Byun S, Seok S, Guo G, Xu HE, Kemper B, Kemper JK. Small heterodimer partner and fibroblast growth factor 19 inhibit expression of NPC1L1 in mouse intestine and cholesterol absorption. Gastroenterology 2019;156(4):1052–1065.

24. Gonzalez FJ. Nuclear receptor control of enterohepatic circulation. Compr. Physiol. 2012;2(October):2811–2828.

25. Kliewer SA, Mangelsdorf DJ. Bile acids as hormones: The FXR-FGF15/19 pathway. Dig Dis 2015;33(3):327–331.

26. Rao A, Haywood J, Craddock AL, Belinsky MG, Kruh GD, Dawson PA. The organic solute transporter -, Ost -Ost, is essential for intestinal bile acid transport and homeostasis. Proc. Natl. Acad. Sci. 2008;105(10):3891–3896.

27. Neimark E, Chen F, Li X, Shneider BL. Bile acid-induced negative feedback regulation of the human ileal bile acid transporter. Hepatology 2004;40(1):149–156.

28. Ballatori N, Christian W V, Wheeler SG, Hammond CL. The heterodimeric organic solute tranporter, OSTα–OSTβ/SLC51: A transporter for steroid-derived molecules. Mol. Aspects Med. 2014;34(0):1–15.

29. Chen F, Ma L, Dawson PA, Sinal CJ, Sehayek E, Gonzalez FJ, Breslow J, Ananthanarayanan M, Shneider BL. Liver receptor homologue-1 mediates species-and cell line-specific bile acid-dependent negative feedback regulation of the apical sodium-dependent bile acid transporter. J. Biol. Chem. 2003;278(22):19909–19916.

30. Davis RA, Miyake JH, Hui TY, Spann NJ. Regulation of cholesterol-7alpha-hydroxylase: BAREly missing a SHP. J. Lipid Res. 2002;43(4):533–543.

31. Frankenberg T, Dawson PA, Haywood J, Shneider BL, Rao A, Chen F. Regulation of the mouse organic solute transporter α-β, Ostα-Ostβ, by bile acids. Am. J. Physiol. Liver Physiol. 2005;290(5):G912–G922.

32. Hegyi P, Maléth J, Walters JR, Hofmann AF, Keely SJ. Guts and gall: Bile acids in regulation of intestinal epithelial function in health and disease. Physiol. Rev. 2018;98(4):1983–2023.

33. Treveil A, Sudhakar P, Matthews ZJ, Wrzesiński T, Jones EJ, Brooks J, Ölbei M, Hautefort I, Hall LJ, Carding SR, Mayer U, Powell PP, Wileman T, Di Palma F, Haerty W, Korcsmáros T. Regulatory network analysis of Paneth cell and goblet cell enriched gut organoids using transcriptomics approaches. Mol. Omi. 2020;16(1):39–58.

34. Zou A, Lehn S, Magee N, Zhang Y, City K. New insights into orphan nuclear receptor SHP in liver cancer. Nucl Recept. Res 2015;2:101162.

35. Garruti G, Wang HH, Bonfrate, Leonilde et al. A pleiotropic role for the orphan nuclear receptor small heterodimer partner in lipid homeostasis and metabolic pathways. J Lipids 2012;2012(604630). doi: 10.1155/2012/304292.

36. Zhou X, Cao L, Jiang C, Xie Y, Cheng X, Krausz KW, Qi Y, Sun L, Shah YM, Gonzalez FJ, Wang G, Hao H. PPARα-UGT axis activation represses intestinal FXR-FGF15 feedback signalling and exacerbates experimental colitis. Nat. Commun. 2014;5(4573):1–15.

37. Halilbasic E, Claudel T, Trauner M. Bile acid transporters and regulatory nuclear receptors in the liver and beyond. J. Hepatol. 2013;58(1):155–168.

38. Li N, Cui Z, Fang F, Lee JY, Ballatori N. Heterodimerization, trafficking and membrane topology of the two proteins, Ostα and Ostβ, that constitute the organic solute and steroid transporter. Biochem. J. 2007;407(3):363–372.

39. Christian W V., Li N, Hinkle PM, Ballatori N. β-Subunit of the Ostα-Ostβ organic solute transporter is required not only for heterodimerization and trafficking but also for function. J. Biol. Chem. 2012;287(25):21233–21243.

40. Li H, Chen F, Shang Q, Pan L, Shneider BL, Chiang JYL, Forman BM, Ananthanarayanan M, Tint GS, Salen G, Xu G. FXR-activating ligands inhibit rabbit ASBT expression via FXR-SHP-FTF cascade. Am. J. Physiol. Liver Physiol. 2004;288(1):G60–G66.

41. Lee Y-K, Goodwin B, Mangelsdorf DJ, Hammer RE, Kliewer SA, Schmidt DR, Zhang Y, Choi M, Peng L, Cummins CL. Liver receptor homolog-1 regulates bile acid homeostasis but is not essential for feedback regulation of bile acid synthesis. Mol. Endocrinol. 2008;22(6):1345–1356.

42. Lan T, Haywood J, Rao A, Dawson PA. Molecular mechanisms of altered bile acid homeostasis in organic solute transporter-alpha knockout mice. Dig. Dis. 2011;29(1):18–22.

43. Jung D, Fantin AC, Scheurer U, Fried M, Kullak-Ublick GA. Human ileal bile acid transporter gene ASBT (SLC10A2) is transactivated by the glucocorticoid receptor. Gut 2004;53(1):78–84.

44. Jung D, Fried M, Kullak-Ublick GA. Human apical sodium-dependent bile salt transporter gene (SLC10A2) is regulated by the peroxisome proliferator-activated receptor α. J. Biol. Chem. 2002;277(34):30559–30566.

45. Ma L, Jüttner M, Kullak-Ublick GA, Eloranta JJ. Regulation of the gene encoding the intestinal bile acid transporter ASBT by the caudal-type homeobox proteins CDX1 and CDX2. Am. J. Physiol. Liver Physiol. 2011;302(1):G123–G133.

46. Hunt MC, Yang YZ, Eggertsen G, Carneheim CM, Gåfvels M, Einarsson C, Alexson SEH. The peroxisome proliferator-activated receptor α (PPARα) regulates bile acid biosynthesis. J. Biol. Chem. 2000;275(37):28947–28953.

47. Shin DJ, Campos JA, Gil G, Osborne TF. PGC-1α Activates CYP7A1 and Bile Acid Biosynthesis. J. Biol. Chem. 2003;278(50):50047–50052.

48. Ikeda I, Metoki K, Yamahira T, Kato M, Inoue N, Nagao K, Yanagita T, Shirakawa H, Komai M. Impact of fasting time on hepatic lipid metabolism in nutritional animal studies. Biosci. Biotechnol. Biochem. 2014;78(9):1584–1591.

49. Jensen TL, Kiersgaard MK, Sørensen DB, Mikkelsen LF. Fasting of mice?: a review. Lab Anim 2013;47(4):225–240.

50. Thompson KL, Applegate TJ. Feed withdrawal alters small-intestinal morphology and mucus of broilers. Poult. Sci. 2006;85(9):1535–1540.

51. Zavarize KC, Sartori JR, Gonzales E, Pezzato AC. Morphological changes of the intestinal mucosa of broilers and layers as affected by fasting before sample collection. Rev. Bras. Cienc. Avic. 2012;14(1):21–25.

52. Zhang Y, Hagedorn CH, Wang L. Role of nuclear receptor SHP in metabolism and cancer ☆. Biochim. Biophys. Acta Mol Basis Dis 2011;1812(8):893–908.

