## Supplemental Figures and Table for "Deletion of intestinal SHP impairs short-term response to cholic acid challenge in mice"

**Supplementary Figure 6. Method for tissue clearing using CUBIC.** (A) Flow chart showing the process for tissue clearing and staining using CUBIC. (B) Before and after images of intestinal tissue using CUBIC.

**Supplementary Figure 7. Intestinal goblet cell counts were maintained during chronic CA diet.** (A-E) Histomorphological analysis using alcian blue staining revealed no statistical difference in the number of goblet cells and crypt depth but there was increase in villi length in *IShpKO* mice after chronic CA diet. Data are represented as mean ±SD, *n*=3-5. Unpaired t-test. (F-J) The expression of *Pparα* and *Ugt* genes regulating glucuronidation of metabolic bile acid for elimination were upregulated in the ileum. Data are represented as mean ±SD, *n*=3-6 and analyzed with a two-way ANOVA, Bonferroni's test.

### Supplemental Figures and Table

### Supplementary Figure 1

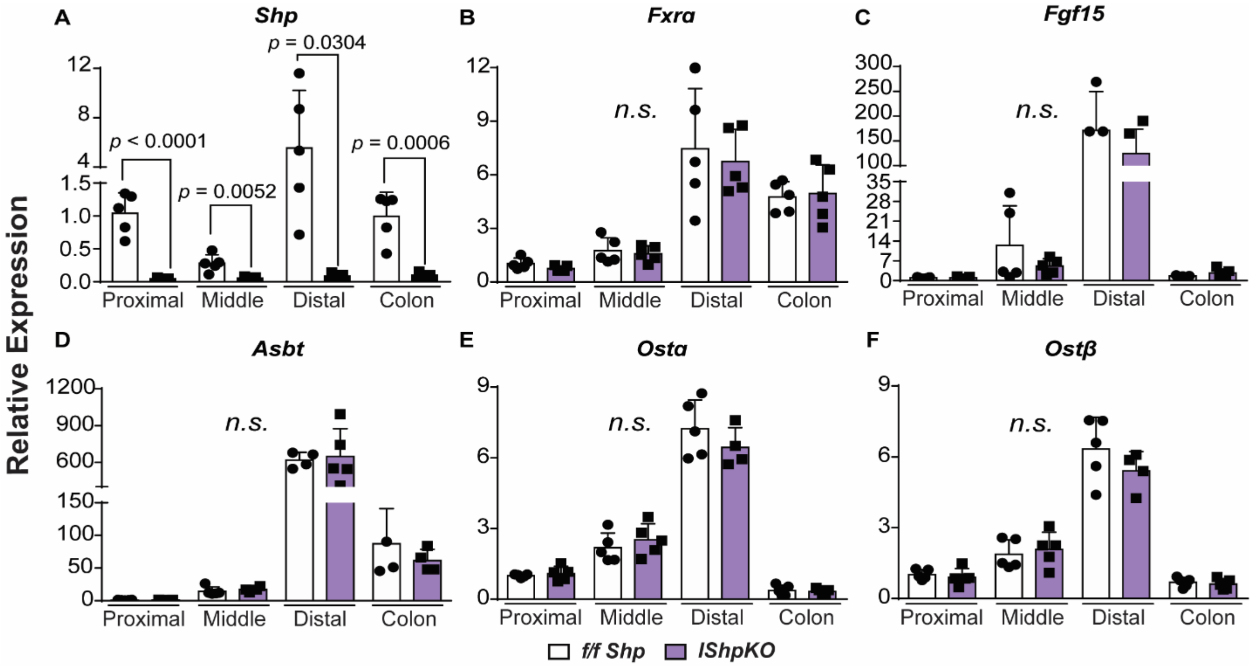

### Supplementary Figure 2

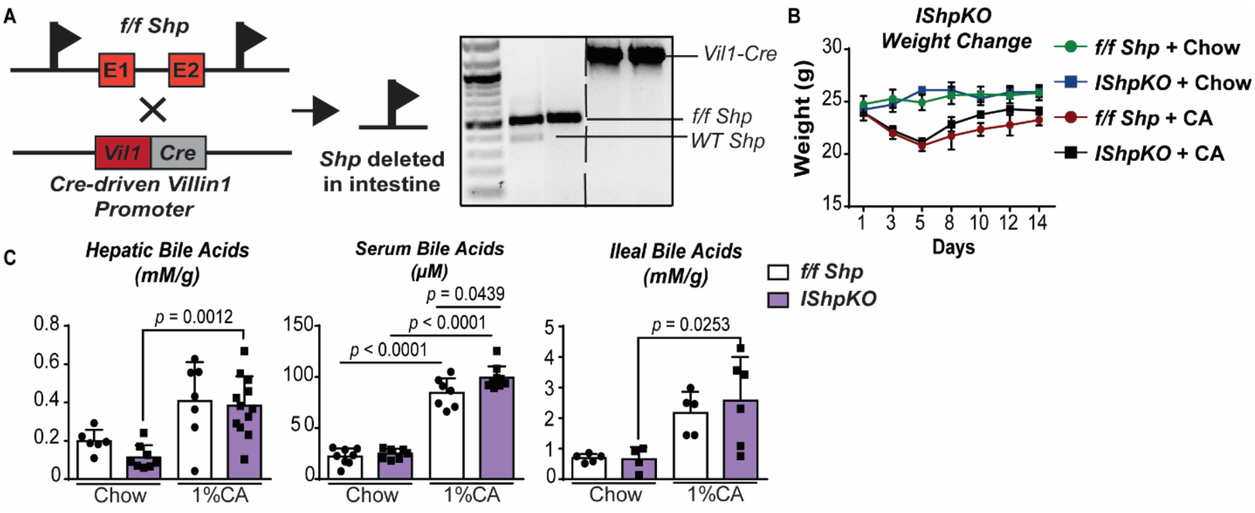

### Supplementary Figure 3

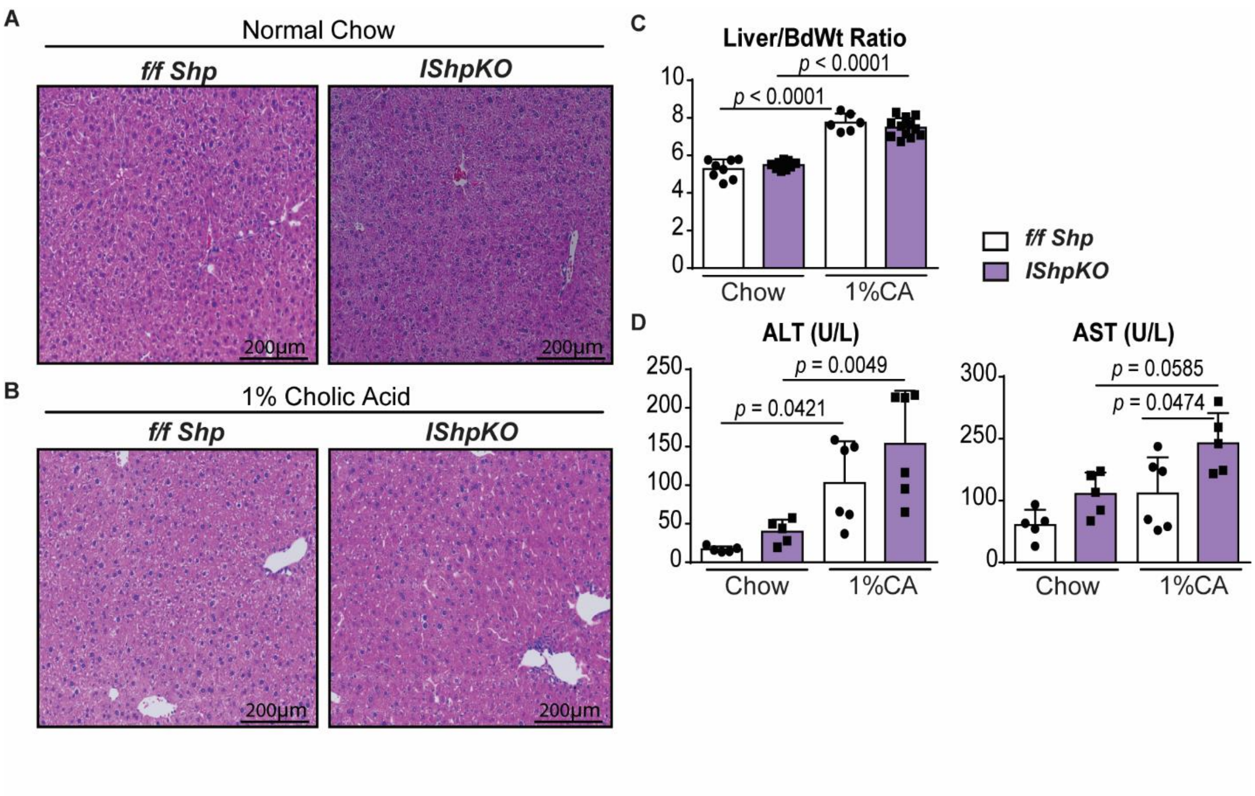

### Supplementary Figure 4

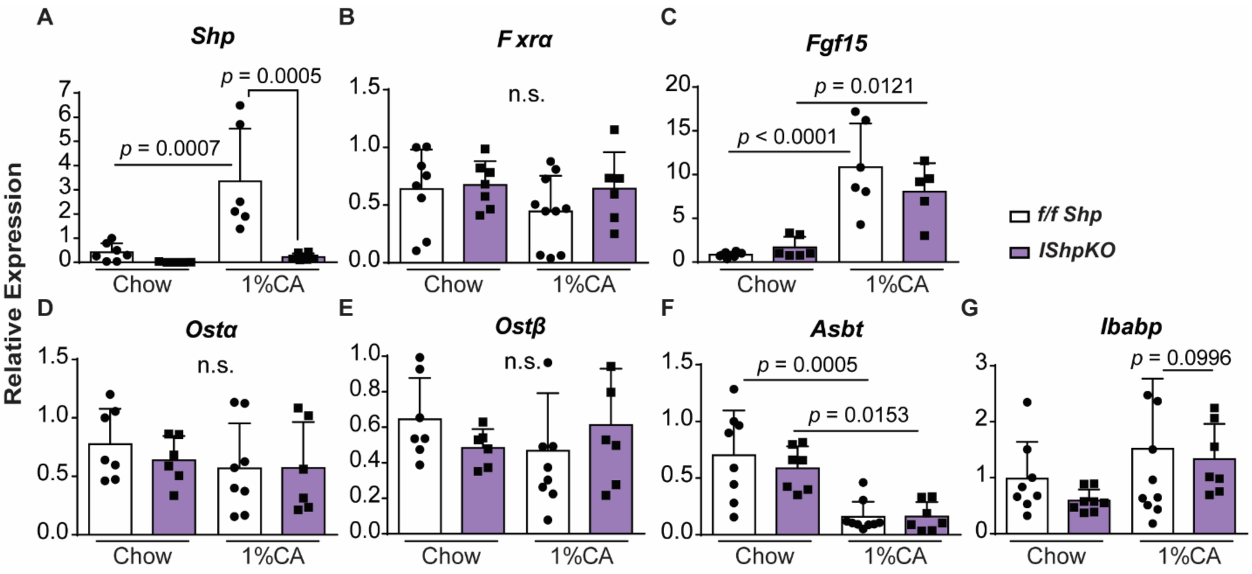

### Supplementary Figure 5

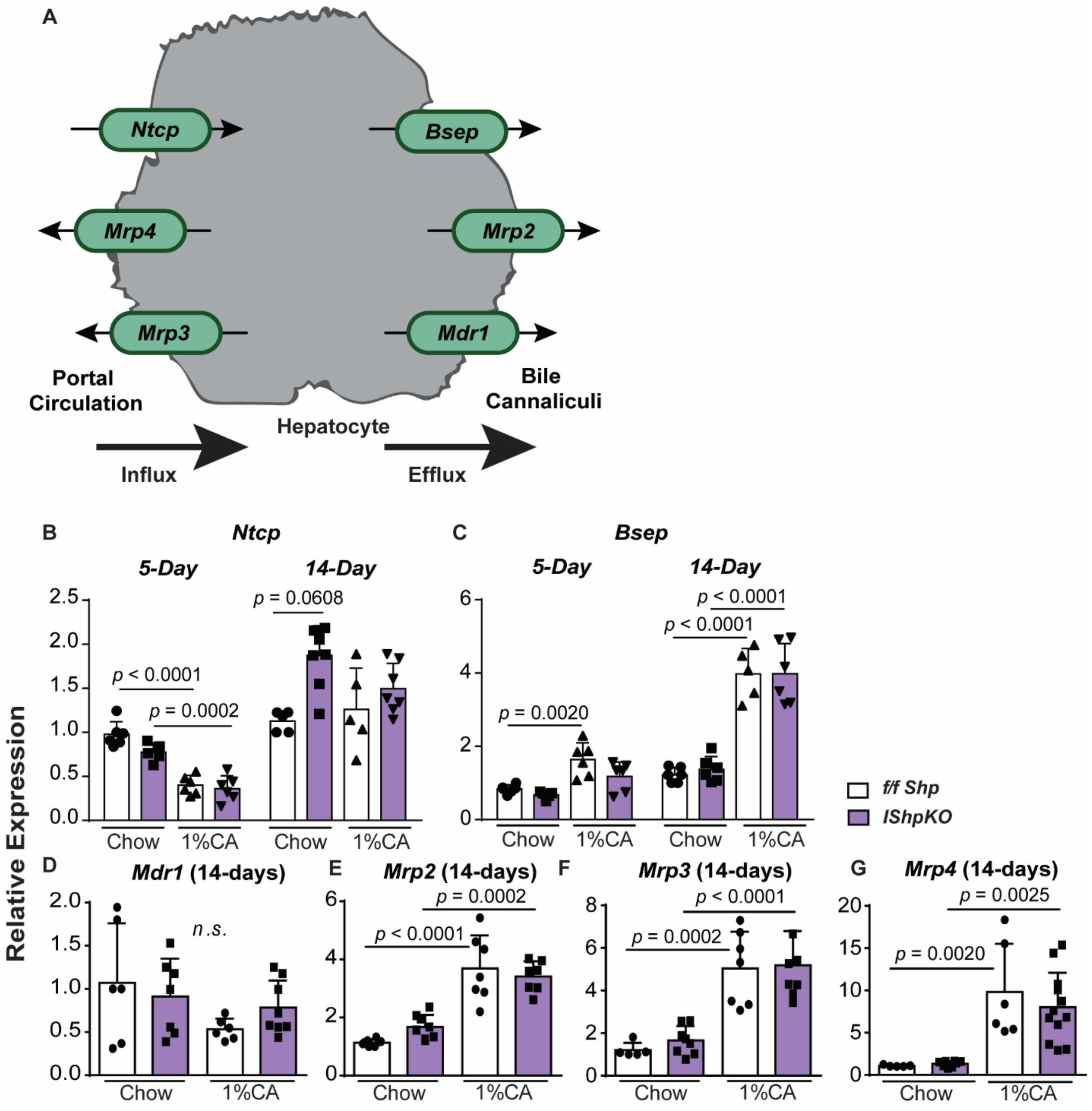

**Supplementary Figure 6**

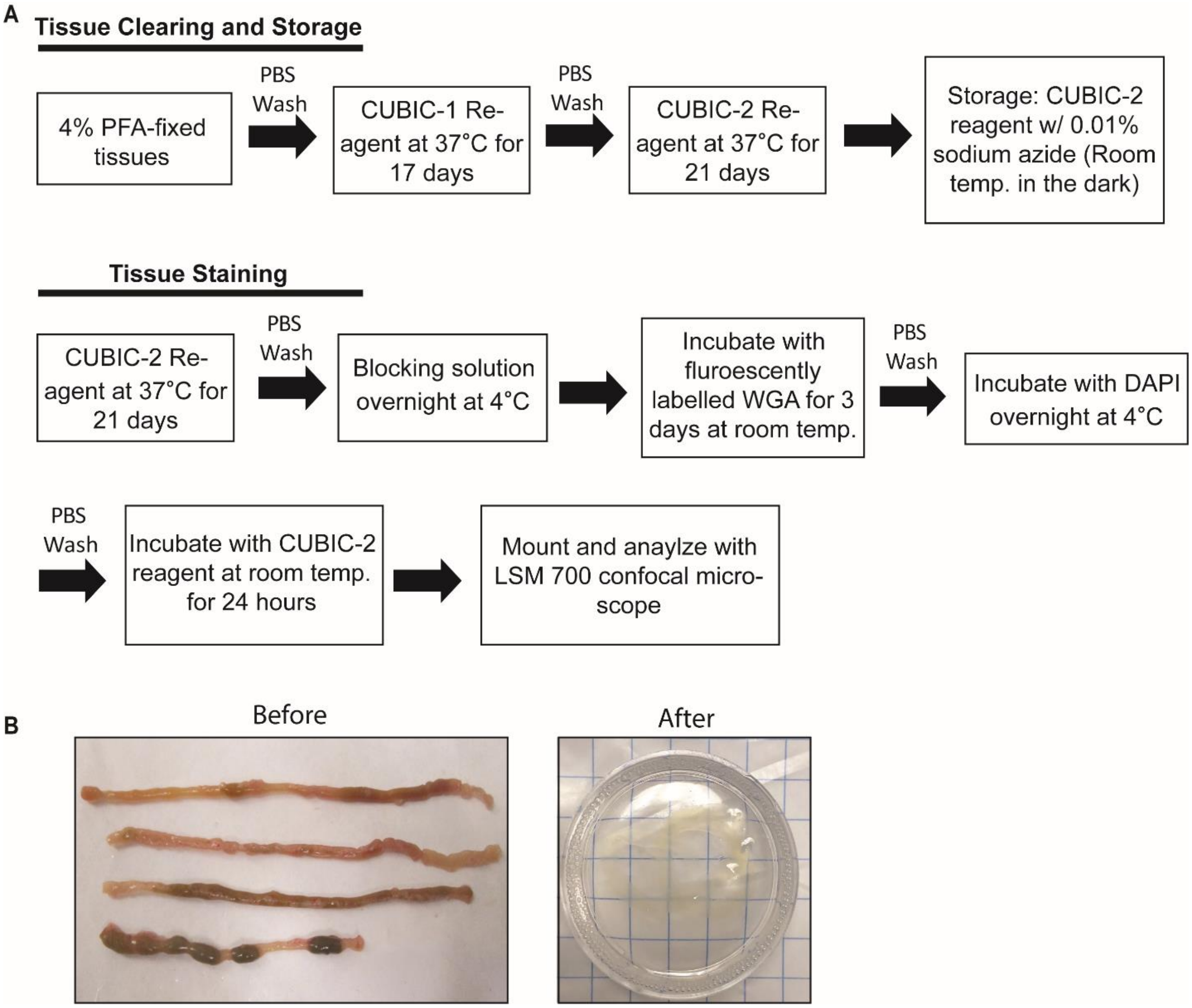

### Supplementary Figure 7

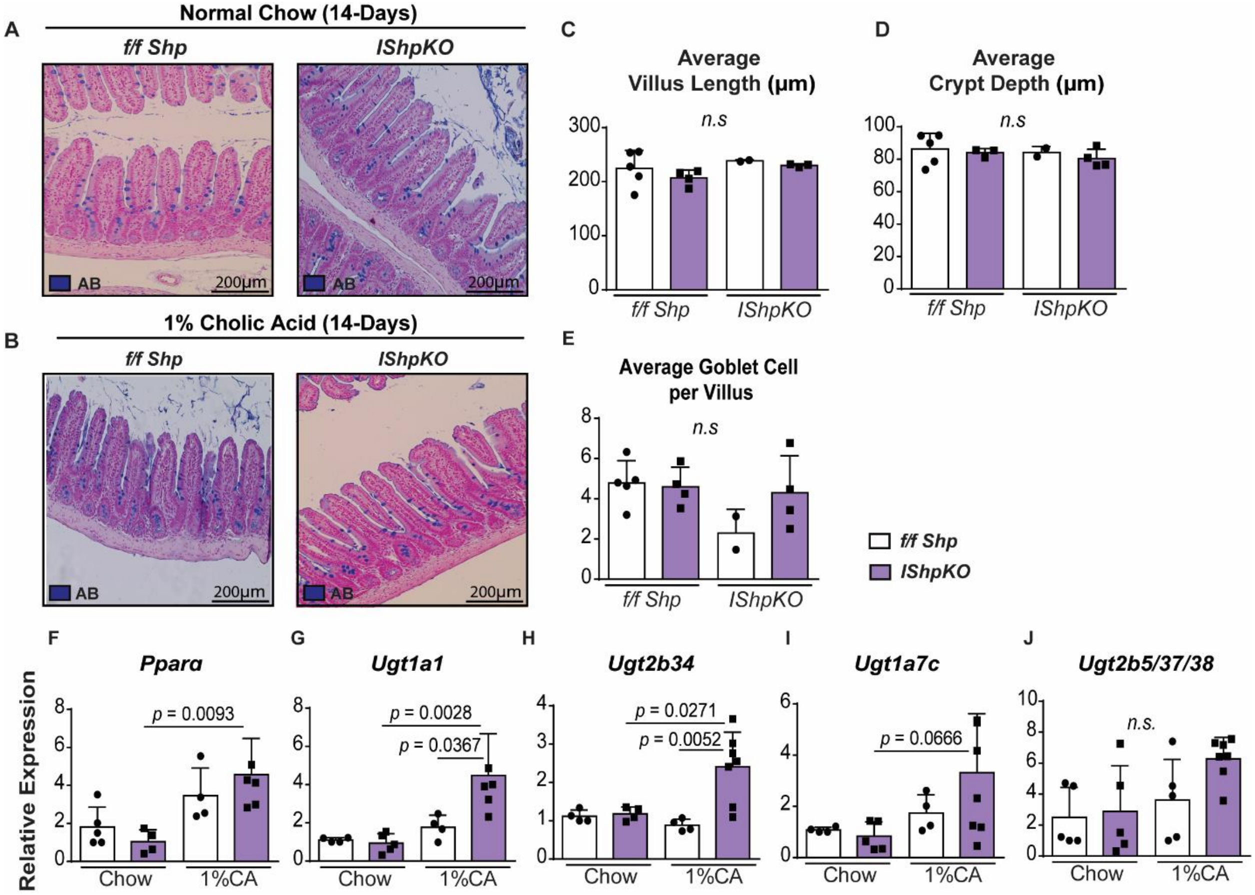

**Supplemental Table 1. qRT-PCR Primers Used.**

| Gene | Forward (5' → 3') | Reverse (5' → 3') |
|---|---|---|
| *36b4/Rplp0* | AGATGCAGCAGATCCGCAT | GTTCTTGCCCATCAGCACC |
| *Gapdh* | AACTTTGGCATTGTGGAAGG | ACACATTGGGGGTAGGAACA |
| *Shp/Nrb02* | CGATCCTCTTCAACCCAGATG | AGGGCTCCAAGACTTCACACA |
| *Fxrα/Nr1h4* | ACAGCTAATGAGGACGACAG | GATTTCCTGAGGCATTCTCTG |
| *Fgf15* | CGCTACTCGGAGGAAGACTG | TTTGAGGGTTTCTGGTCCTG |
| *Asbt/Slc10a2* | GGAACTGGCTCCAATATCCTG | GTTCCCGAGTCAACCCACAT |
| *Ostα/Slc51a* | ATGCATCTGGGTGAACAGAA | GAGTAGGGAGGTGAGCAA |
| *Ostβ/Slc51b* | GACCACAGTGCAGAGAAAGC | CTTGTCATGACCACCAGGAC |
| *Ibabp/Fabp6* | GGTCTTCCAGGAGACGTGAT | ACATTCTTTGCCAATGGTGA |
| *Ntcp/Slc10a1* | CTCAGCGTCATTCTGGTAGTT | CCAGAAGTGAGCCTTGATCTT |
| *Bsep/Abcb11* | CCAGAGGCAGCTATCAGGAC | CACACAAAGCCCCTACCAGT |
| *Cyp7a1* | CCTAAGAGCAAAGCAAAGGAAAC | CTTTGTGGTATGACAGGGAGTT |
| *Cyp8b1* | TTTCTGAGGGAGCAAGGAATAG | GGAATAAGAGGACCCAGAAACA |
| *Pparα* | ACAAGGCCTCAGGGTACCA | GCCGAAAGAAGCCCTTACAG |
| *Ugt1a1* | ATGGCTTTCTTCTCCGGAAT | TCAGAAAAAGCCCCTATCCC |
| *Ugt1a7c* | TGCAATGGAGTTCCGATGGT | CTGGAGAGGCGCATGATGTT |
| *Ugt2b34* | GGAGAATGCCATGCGGTTAT | CTGCCACACGAAGATGCTTG |
| *Ugt2b5/37/38* | TGGCCGATGGAATTCAGTC | GTTTCAAACTTAAGGCCAGGTG |
